# Using agent-based models to predict pollen deposition in a dioecious crop

**DOI:** 10.1101/2022.07.28.501917

**Authors:** Melissa A. Broussard, Mateusz Jochym, Nathan Tomer, Linley Jesson, Allison K. Shaw, David W. Crowder, Nilsa A. Bosque-Pérez, Jing Li, Angela Peace, Dilini Fonseka, Brad Howlett, David Pattemore

## Abstract

Pollination involves complex interactions between plants and pollinators, and variation in plant or pollinator biology can lead to variability in pollination services that are difficult to predict. Models that effectively predict pollination services could enhance the ability to conserve plant-pollinator mutualisms in natural systems and increase crop yields in managed systems. However, while most pollination models have focused either on effects of plant or pollination biology, few models have integrated plant-pollinator interactions. Moreover, crop management causes variation in plant-pollinator interactions and pollination services, but management is rarely considered in pollination models. Here we used extensive datasets for kiwifruit (*Actinidia chinensis* var. *deliciosa*) to develop an agent-based model to track insect-provided pollination services with variation in crop cultivars, pollinator traits, and orchard layouts. This allowed us to predict pollination outcomes in a dioecious crop under a range of management scenarios. Our sensitivity analysis indicated that flower density and the proportion of female flowers are the most important factors in successful pollination, both of which growers control via cultivar selection and cultural management practices. Our analysis also indicated that economically viable pollination services and crop yields are attained with ∼60% female flowers and a peak foraging activity of 6 to 8 bees per 1,000 open flowers with diminishing returns for additional pollinators. The quality of pollination service varied across simulated orchard layouts, highlighting the potential use of this model as a framework to screen novel orchard configurations. More broadly, linking complex plant and pollinator interactions in pollination models can help identify factors that may improve crop yields and provide a framework for identifying factors important to pollination in natural ecosystems.

**HIGHLIGHTS:**

- We develop a model using extensive empirical datasets to predict pollen deposition based on the interactions between flowers and pollinators in a dioecious crop system
- We conducted a thorough sensitivity analysis, and analysis of the effect of stochastic variance between model runs, which can be used to inform future design of stochastic agent-based models
- Our model effectively predicted the outcomes of varying management regimes of orchard layouts and pollinator introductions on pollination in a dioecious crop
- Our model can be extended for other functionally dioecious crops or plant communities where managers want to understand how their decisions impact pollination

## 1 Introduction

Insect-mediated pollination is a critical ecosystem service that is responsible for the production of the majority of fresh fruit and vegetable crops globally (Klein et al. 2007). Crop pollination deficits are common across a broad range of crops grown within agroecosystems worldwide (Garibaldi et al. 2013), indicating that an improved understanding and management of pollination could increase crop yields. Despite the critical importance of pollination for yield, it is a complex process that remains poorly understood, and empirical studies are often only able to assess the effect of one or two variables at a time (e.g. (Carvalheiro et al. 2010; Holzschuh et al. 2011; King et al. 2013; Miñarro & Twizell 2015). This is problematic because multiple interacting factors from plants, pollinators, weather, farm management practices and the landscape context affect pollination deficits and crop yields, and empirical studies may fail to effectively capture key factors.

For functionally dioecious crops, the timing of flowering is also critical, especially the overlap between pollen donor (i.e. ‘male’) flowers and the receptive ‘female’ flowers, as this overlap governs the window of time in which pollination can occur. At present, growers have limited options for managing pollination, principally the short-term practice of bringing in managed bee hives each season and the significantly more costly long-term investment of establishing planting regimes of cultivar combinations, or male and female plants, which may last for a period of time from one year to more than a decade (Delaplane & Mayer 2000). Unfortunately, there is little robust information available to provide recommendations to growers about hive stocking rates or planting regimes, so inappropriate stocking rates and field designs are likely to contribute to yield deficits (Rollin & Garibaldi 2019).

Modelling has greatly assisted predictive management in other areas of agriculture that involve multiple interacting factors that are difficult to adequately control and replicate in empirical field studies (Sinclair & Horie 1989; Bonhomme 2000; Valipour et al. 2015). Pollination is a critical, but challenging agricultural service which has received attention from a number of modelers. However, most models of pollination have focused either on the crop with limited consideration of pollinator behavior and dynamics (e.g. Brain & Landsberg 1981; Degrandi-Hoffman et al. 1987; Lescourret et al. 1988; Lescourret et al. 1998) or on the dynamics of pollinator abundance across a landscape without considering their interaction with flowers (Lonsdorf et al. 2009). There is a need for models that examine both factors simultaneously (Qu & Drummond 2018). Qu & Drummond (2018) concluded that agent based models (ABMs) were a useful simulation tool to evaluate the influence of a wide variety of plant, pollinator and environmental factors that affect pollination efficiency and yield.

ABMs have been used to explore several facets of pollination, including pollination efficiency (Qu et al. 2013), the effects of bee density-dependent preferences on pollination (Rands & Whitney 2010), crop planting requirements to sustain a pollinator species (Dumont et al. 2018), interactions between pesticides, climate, biodiversity and pathogens and their impact on bee health and navigation (Becher et al. 2014; Betti et al. 2017), and the influence of landscape features on bee behaviour (Rands 2014), and the relative pollination efficiencies of different bee taxa (honey bees, bumble bees and solitary bees) including the influence of foraging distance from nest sites (Qu & Drummond 2018). However, we are unaware of examples where ABMs have been used to predict optimal field design and bee densities in relation to functionally dioecious crops.

Our innovation was to develop a spatially-explicit ABM that captured the dynamics of pollination in a dioecious crop, allowing for a fine-grained ability to test the effects of manipulating key management actions. Additionally, we used a two-tiered approach to ensure a) adequate model replication at each parameter combination and b) that enough Latin hypercube samples were taken, reducing the impact of stochasticity on our analysis.

Here, we tested the impact of three broad classes of factors on pollination success:

1. The effects of plant-biology-focused metrics, including flower density, the ratio of female to male flowers, phenology, and pollen production.
2. The impact of pollinator-biology-focused metrics, including bee density, pollen removal efficiency, grooming efficiency, and floral sex preference.
3. The impact of different orchard layouts on the relative importance of plant and pollinator biology on crop pollination.

Through this integrated approach our model provided novel insight into how plant and pollinator biology interact with management factors to affect pollination services in a dioecious crop.

## 2 Methods

### 2.1 Study System

Kiwifruit (*Actinidia chinensis* var. *deliciosa*, A Chev) is a dioecious, high-value crop for which insect pollination is essential to produce economic yields (Palmer-Jones & Clinch 1974; Schmid 1978; Costa et al. 1993; González et al. 1998; Klein et al. 2007). Neither floral sex produces nectar, and the female flowers produce nonviable pollen as an insect reward (Schmid 1975). The flowers are syncarpous (Vaissière et al. 1990), meaning that pollen deposition can be modelled at the flower rather than the carpel level, as the interconnectivity means that full fruit set can occur from pollen being deposited on any stigma. Extensive datasets on key pollination parameters have been collected over the last three decades, making it possible to use kiwifruit as a model system for other dioecious and functionally dioecious crops.

A number of factors have been shown to affect seed and fruit set rates in kiwifruit, including bee density (Palmer-Jones & Clinch 1974; Matheson 1991), aspects of bee behaviour and movement (Blanchet 1987; Goodwin & Steven 1993), pollinator diversity (Miñarro & Twizell 2015), and the phenology and pollen quantity of male cultivars (González et al. 1994).

The purpose of this model is to create a framework which is capable of predicting pollen deposition, and thereby fruit set, in kiwifruit given an orchard configuration and the density of pollinators per 1000 flowers.

### 2.2 Model Design

Our agent-based model has two agent classes: flowers and pollinators. The model is spatially explicit, with a grid-layout for the orchard landscape which contains a number of stationary flower agents (Figure 1). The final output of the model was the number of pollen grains deposited on each kiwifruit flower during its window of receptivity. Empirical data from the literature were used to parameterize the model (Table 1). Here we use the ODD (Grimm et al. 2010) protocol to briefly describe the model:

**Figure 1:**
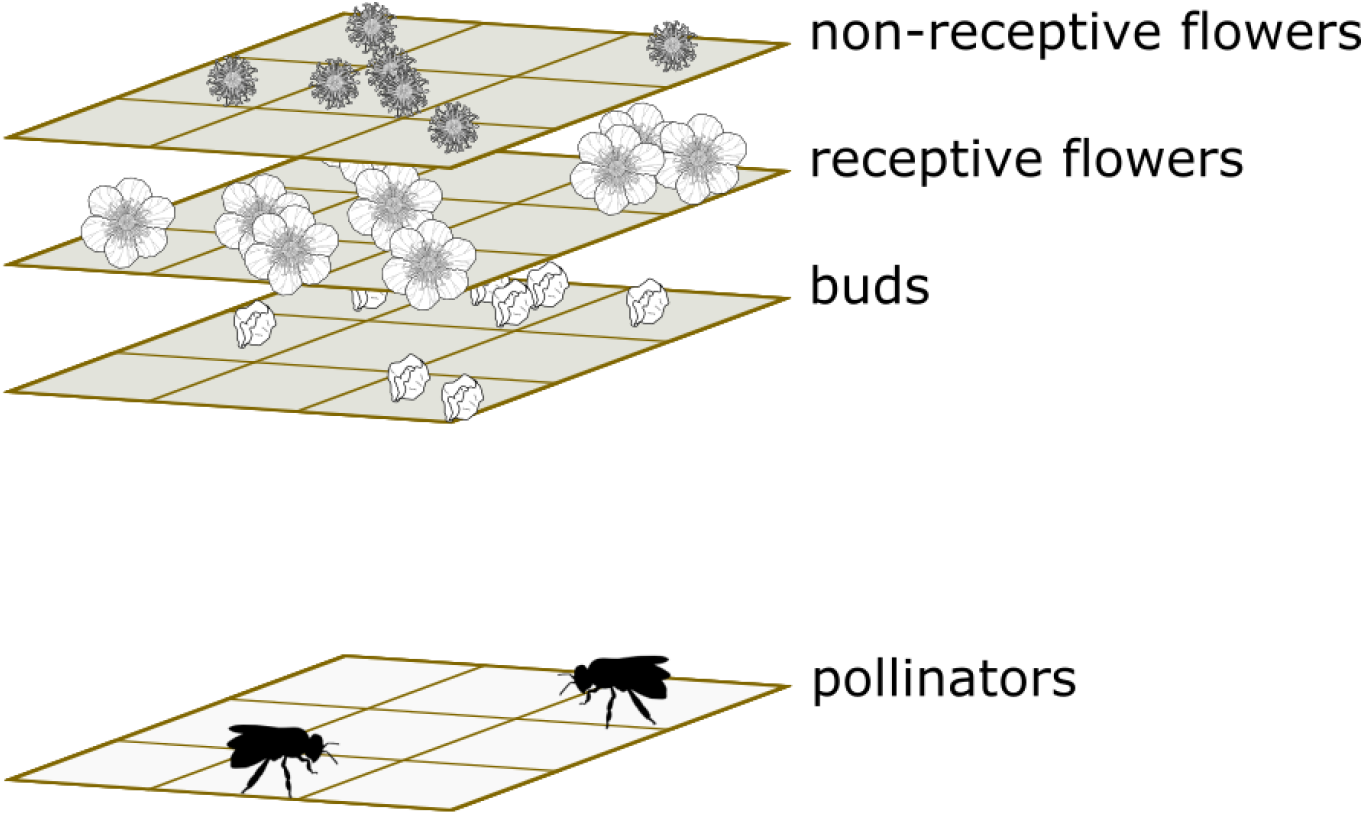
Simplified layout of the landscape and agents in the ABM. The landscape matrix is separated into three layers, with receptive flowers the only ones able to receive visits from pollinators. Pollinator agents can visit any open flower in the tile they are currently occupying.

**Table 1:**
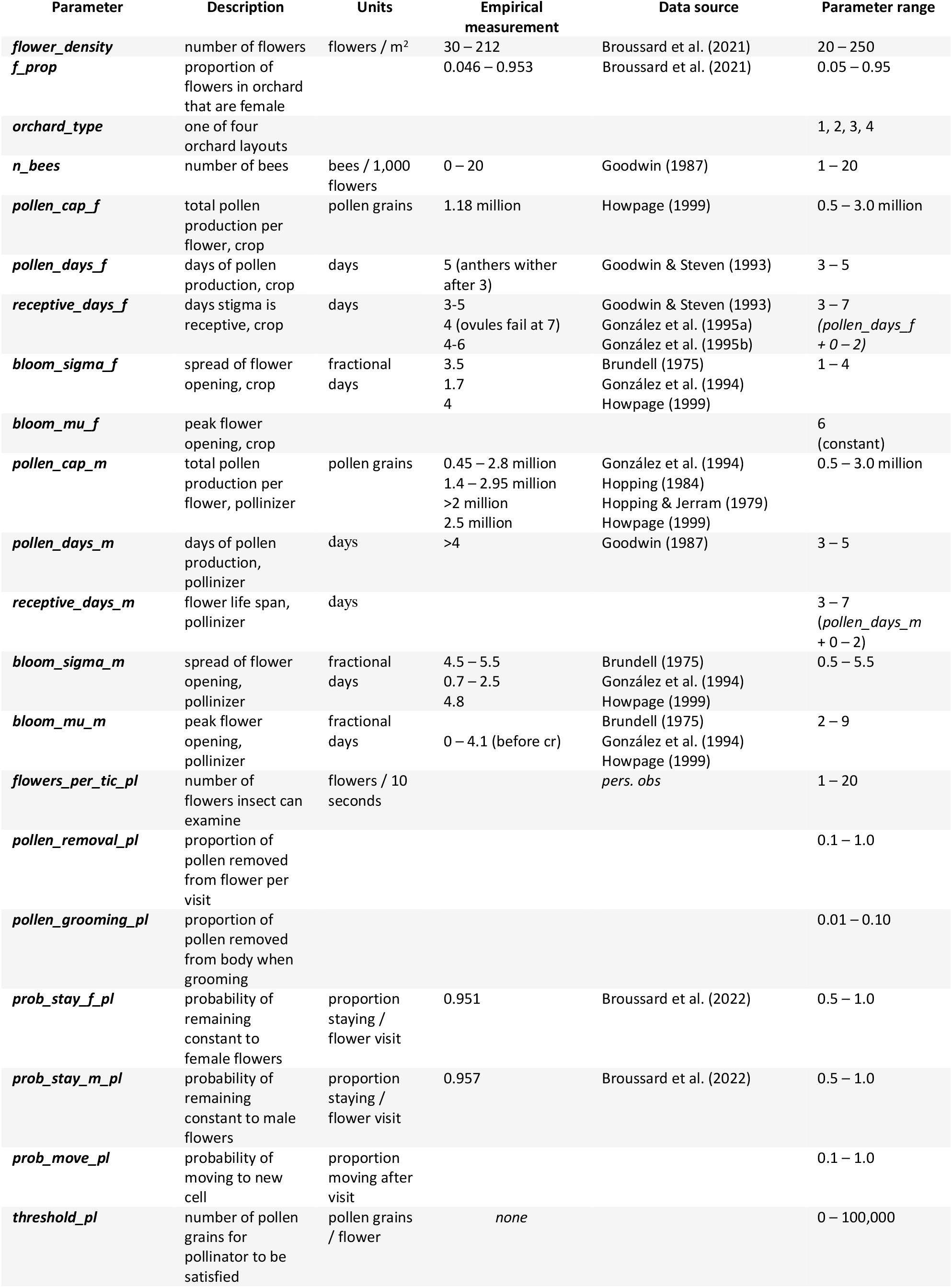
Parameters used in the agent-based model of kiwifruit pollination, empirical data, and the parameter ranges explored during sensitivity analysis.

#### 2.2.1 Agents and the orchard landscape

*The model simulated different configurations of flowers and pollinators in a landscape. Flowers were characterized by: (i) their sex, (ii) the number of pollen grains produced, (iii) the time at which they are scheduled to open, (iv) the number of days pollen is produced, and, (v) for female flowers, the number of days the flower remains receptive and (vi) the number of pollen grains received. Based on empirical observations, flower opening time was modelled as a normal distribution observations (Brundell 1975; González et al. 1994; Howpage 1999) defined by σ = bloom_sigma_f, μ = bloom_mu_f for female flowers, and σ = bloom_sigma_m, μ = bloom_mu_m for male flowers (*

Table 1). Each flower was assigned an opening time at random from within the relevant distribution. Pollinator agents were assigned species-level attributes shown to be important in the literature: (i) search rate (Gilman et al. 2012), (ii) proportion of pollen removed in a foraging bout (Harder & Thomson 1989), (iii) proportion of pollen groomed off of the body after a foraging bout, (iv) preference for male or female flowers (Goodwin & Steven 1993), (v) the probability of switching from male to female and vice versa (Delaplane & Mayer 2000), and (vi) the probability of moving to a different area in the orchard (Wratten et al. 2003; Brosi et al. 2008). We also included variables to represent bee satisfaction, as numerous papers have indicated that bees will judge the quality of the foraging patch by visiting subsequent flowers, and, if the level of resources is low, bees will move to another patch (Heinrich 1979; Ott et al. 1985; Harder 1990). These variables are: (i) a pollen threshold below which the bee is unsatisfied and (ii) the number of successive unsuccessful flower visits before the bee will be unsatisfied enough to move from the local patch. Pollinators also track (i) how much pollen is on its body, available for pollination, (ii) how much pollen has been removed (e.g., into the pollen basket), (iii) its current flower sex preference, and two ‘dissatisfaction’ variables: (iv) one which is binary satisfied/unsatisfied based on the most recent flower visit, and (v) a cumulative measure of dissatisfaction.

Pollinator abundance was fixed as a number per 1,000 open kiwifruit flowers in order to both align with historical measures in the literature (Palmer-Jones & Clinch 1974; Clinch 1984; Goodwin 1987; Goodwin et al. 2017) and because it is a simple measure for orchardists to take in the field. The number of pollinators per 1,000 flowers was set between 1 and 20, as this is the range observed in many years of work in orchards (Goodwin 1987, Goodwin pers comm).

The orchard landscape was simulated by 8 × 8m grid cells with different layers for unopened flower buds, opened (receptive) flowers, opened (non-receptive) flowers, and pollinators (Figure 1). For our model runs, we chose a grid cell size of 1m x 1m, large enough to encompass numerous flowers, but small enough to include spatial variation in flower sex and density observed in kiwifruit orchards. Within each grid cell, agents can interact in a spatially implicit fashion. The landscape-level variables explored were (i) the number of pollinators per 1,000 open flowers, (ii) the orchard-level flower density per square meter, (iii) the ratio of male flowers to female flowers, and (iv) the planting layout of the orchard. We used four hypothetical orchard layouts to test our model: (i) the theoretically optimal ‘uniform’ layout, where there is no spatial difference in flower density, (ii) a ‘checkerboard’ layout, where landscape tiles alternate between male and female, (iii) the ‘stripmale’ layout, which is commonly used in New Zealand, and (iv) a less-than-optimal ‘point’ layout, where all male flowers are in a small portion of the orchard (Figure 2). To simulate a larger world than the 8 × 8m grid, bees were able to ‘wrap-around’ the landscape torus, eliminating edge effects. Each timestep represents 10 seconds, with the model being run over the typical period of kiwifruit flowering (14 days; 120,960 timesteps). This timestep allowed us to model pollinator behavior at a fine scale over the flowering period and avoided the computational issues around smaller timesteps.

**Figure 2:**
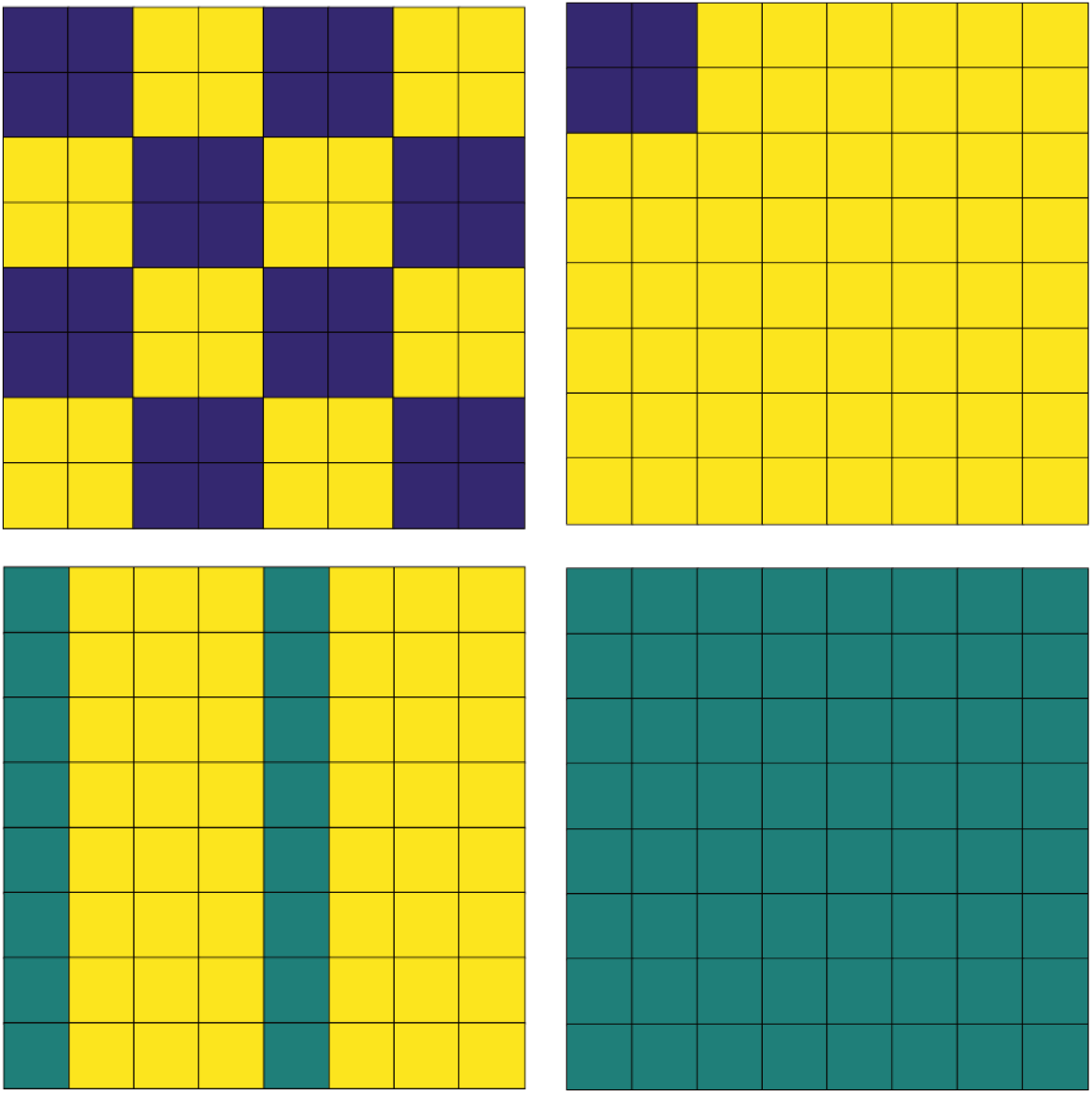
Orchard layouts used in the kiwifruit pollination ABM. Dark blue cells contain only male flowers, yellow cells contain only female flowers. Green cells contain both male and female flowers. The names of the orchards used in this paper are ‘checkerboard’ (top left), ‘point’ (top right), ‘stripmale’ (bottom left; a common layout used by orchardists), and ‘uniform’ (bottom right).

**Figure 2:**
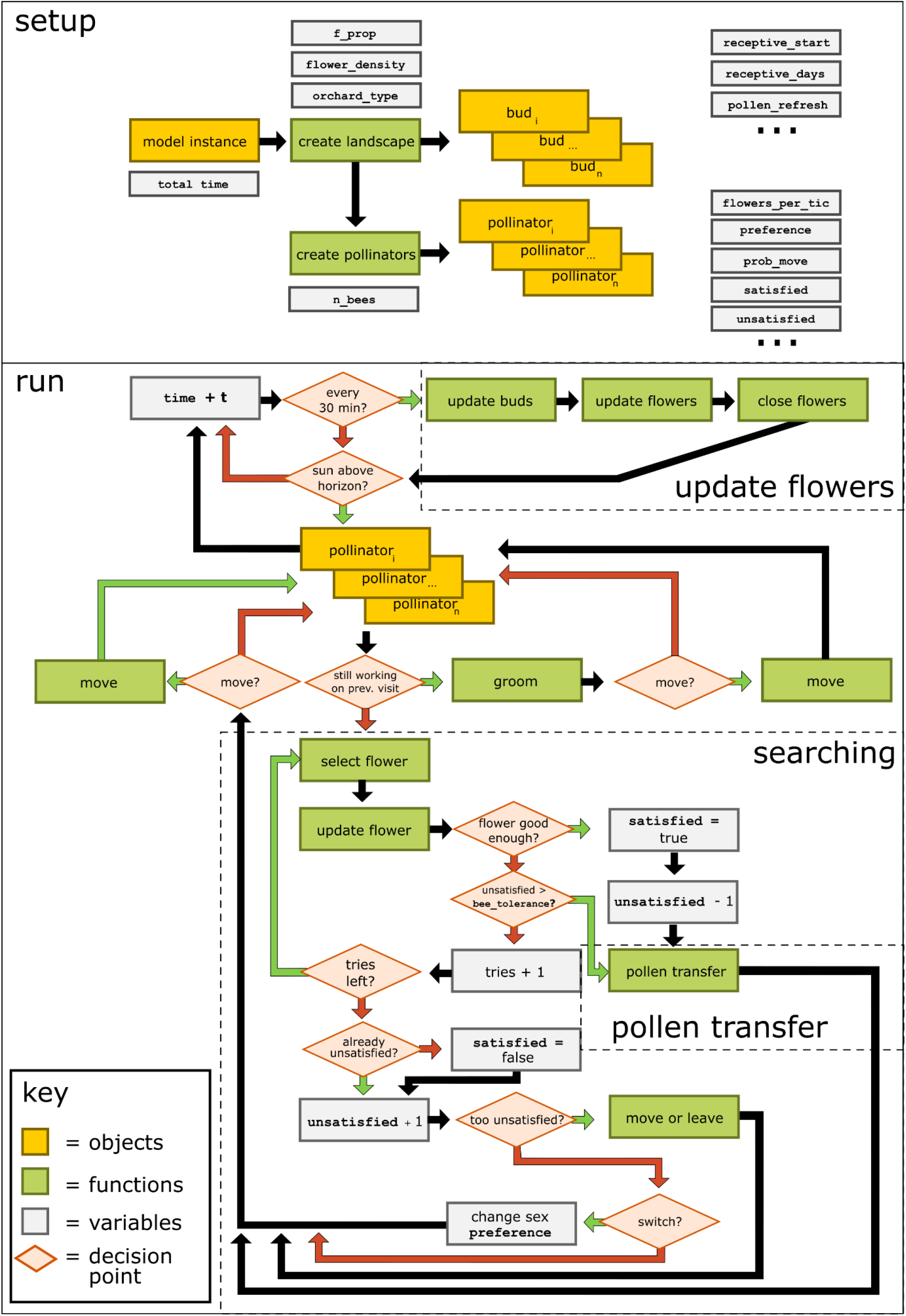
Model logic for our representation of kiwifruit pollination. Yellow boxes indicate agents, green objects are actions, grey boxes are agent variables, and red diamonds are yes/no questions which affect the path of model logic. Red arrows indicate a “no” evaluation, whereas green arrows indicate a “yes” evaluation.

#### 2.2.2 Processes

##### 2.2.2.1 Flower updating

To reduce computing time, flowers are not updated every timestep, but once every 30 minutes of simulated time as well as at the time they are interacted with by a pollinator (Figure 3). During this update, every flower in the bud and receptive flower layer are checked against the current timestep. Buds which are scheduled to open are moved to the receptive flowers, and open flowers which have been open longer than their receptivity window are moved to the non-receptive flower layer. Pollen production for open flowers is updated at this time—flowers produce a flat number of pollen grains per timestep equal to their total pollen production over their receptive window.

**Figure 3:**
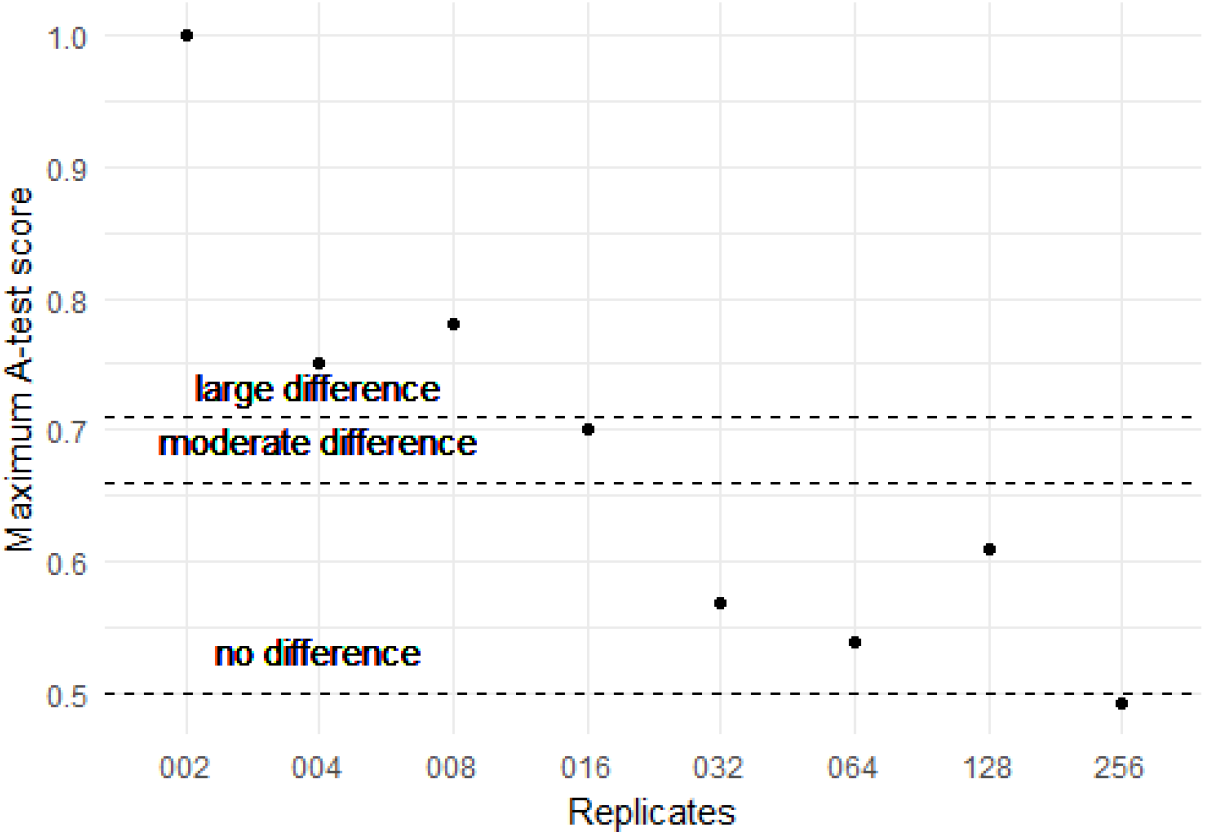
A-test scores for data distributions for different numbers of replicates of the agent-based model. A score closer to 0.5 indicates that there was no difference between the mean of the first run and each of the subsequent runs out of 20 total for each replicate value. Significance values are from Vargha & Delaney (2000).

##### 2.2.2.2 Pollinator foraging

Pollinator agents are able to forage when the sun is above the horizon. This condition is evaluated by custom C code which calculates the sun’s angle given the time and the orchard’s GPS coordinates using the equations described by Blanco-Muriel et al. (2001). For our model runs, the location was set to be Te Puke, a center of kiwifruit production in New Zealand, and the time was set to be the beginning of kiwifruit flowering in the same year our reference data were collected (November 2018).

Every daylight timestep, each pollinator agent attempts to find a suitable flower within its current grid cell. A number of flowers up to the maximum it can examine are selected at random and assessed as to whether the pollen is above the current threshold. If so, the pollinator will attempt to pollinate the flower, where it deposits a proportion of the pollen on its body to the selected flower, as well as removing pollen from the anthers (Figure 4). If the bee is unable to find a suitable flower, it becomes unsatisfied, and adds one to its unsatisfied score (Figure 3). There is a chance every timestep for the bee to switch floral sex preference or move to one of the 8 neighboring grid cells.

**Figure 4:**
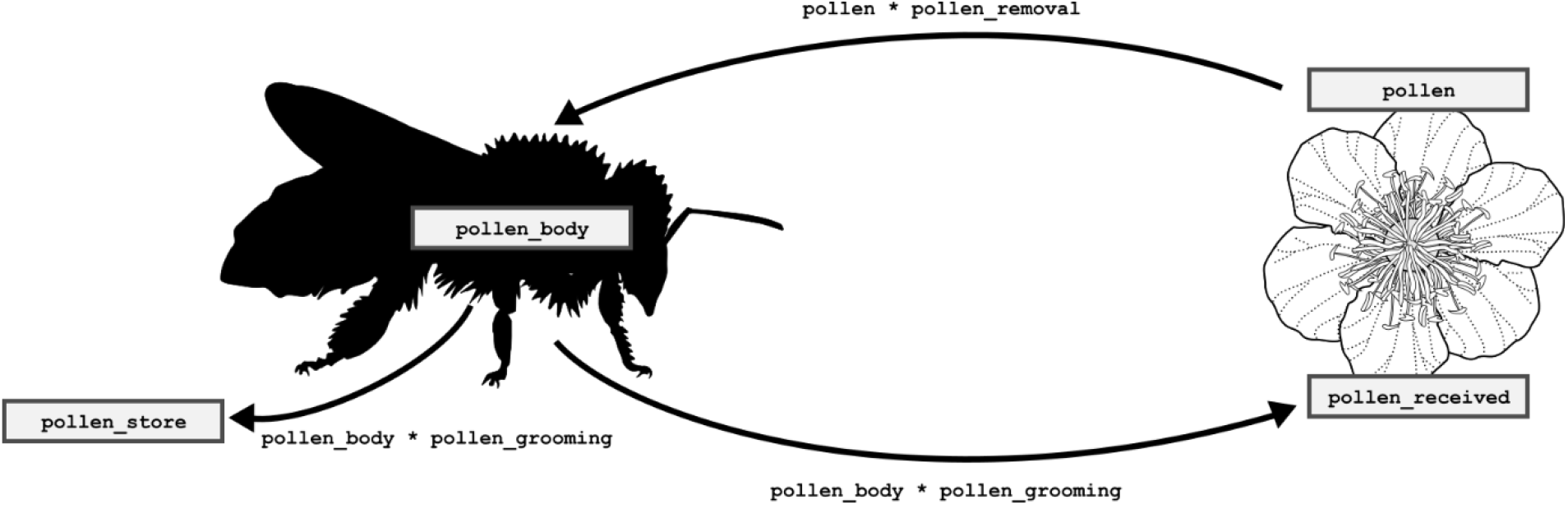
Diagram representing the pollen transfer functions. When a bee visits a flower, it receives pollen in proportion to its pollen removal ability (a proportion between 0.1 and 0.9). The insect deposits pollen on the flower according to its grooming ability (a proportion between 0.01 and 0.1). The bee then grooms itself, moving pollen from its body to its pollen stores, no longer available for transfer. If a bee receives more than three times its pollen threshold, it will take an additional 1-3 timesteps to groom itself, proportional to the pollen received.

**Figure 4:**
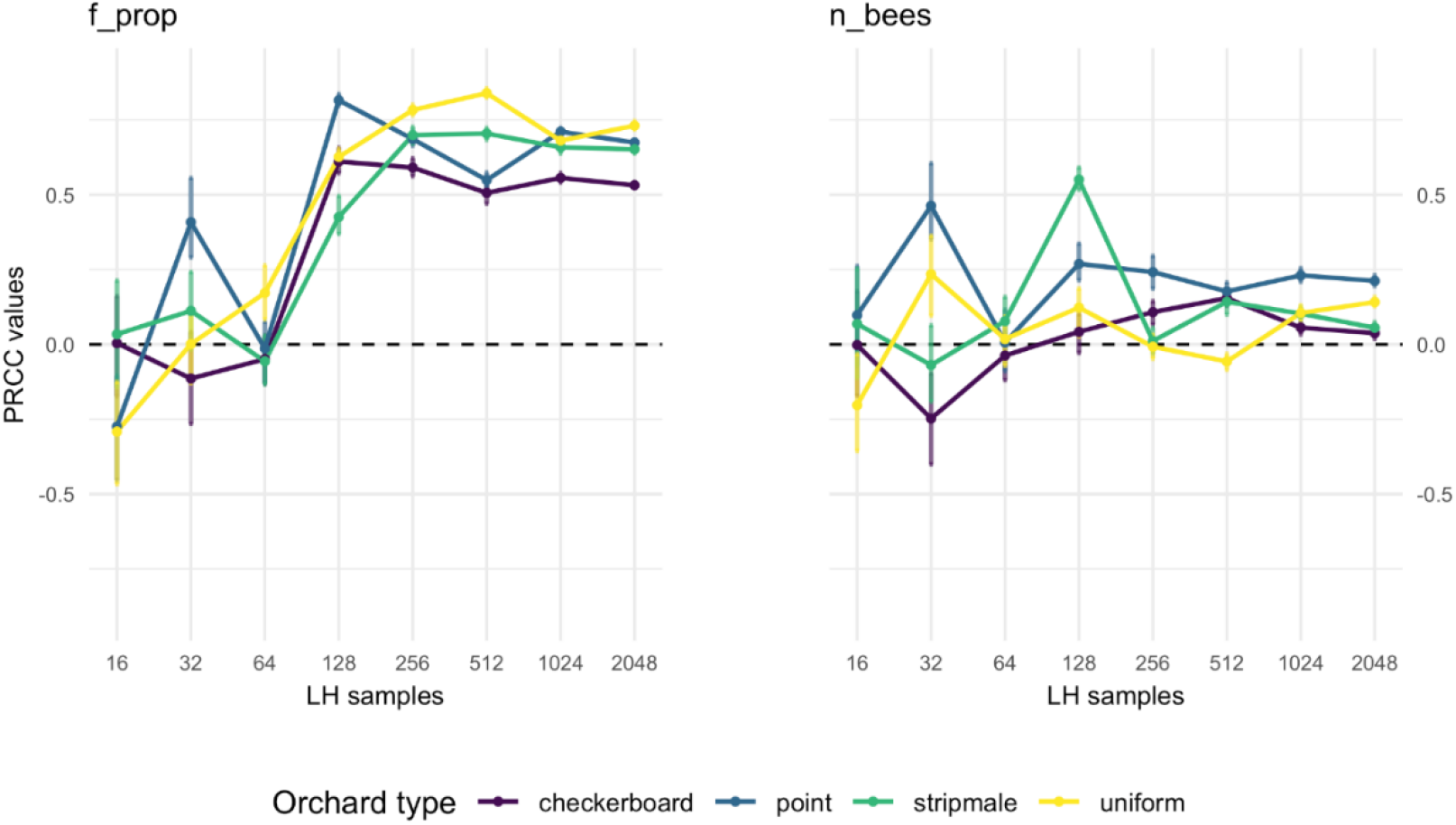
Results of PRCC-based sensitivity analysis for the proportion of female flowers (f_prop) and the number of bees per 1,000 open flowers (n_bees) across the 8 Latin hypercube sampling combinations tested. Results appear to stabilize after 256-512 LHS combinations.

#### 2.2.3 Technical specifications

The model was written in a combination of Python, Cython and C. We initially explored using Python exclusively but, due to the number of agents (961 – 12,240), fine timestep, and long model duration, speed improvements from Cython and C were desirable. We ran the model in a high performance computing environment running on CentOS Linux 7, with OpenLava 3.2.0 load scheduling facility. Model runs were automated using a custom R script (R Core Team 2017) with a foreach loop from the R package *foreach* (Microsoft & Weston 2017).

### 2.3 Sensitivity Analysis

To evaluate the importance of each variable, and explore the boundaries of the parameter space which result in sufficient pollen deposition, we used Latin hypercube sampling (LHS). This method, first described by McKay et al. (1979) allows for a significant reduction in the number of model runs in comparison to sampling each unique combination of parameters. A full-factorial exploration of all parameters (Table 1) would have entailed over 100 quadrillion model runs, taking more than 100 billion years of computation time. Using LHS allowed us to examine the parameter space in 500-2000 runs, taking 15-30 min, a reduction of computing time by 10^15^-fold. The LHS matrices were generated in R using the function *maximinLHS* from the package *lhs* (Carnell 2019) and scaled according to the ranges for each variable analysis (Table 1).

To determine the number of times the model should be run with each parameter combination, we used the approach described in Read et al. (2012), and ran the model with the base set of parameters (observed means) with different numbers of replicates to determine the point at which the distribution of results is consistent. For our model, we ran 20 tests for each of the following number of replicates: 2, 4, 8, 16, 32, 64, 128, and 264. We then used a Vargha-Delaney A-Test (Vargha & Delaney 2000) to determine if distributions varied within the 20 tests with the function *atest* in R package *spartan* (Alden et al. 2015).

With the determined number of replicates, we then ran our model with different numbers of parameter combinations using LHS—an approach used in some other studies of stochastic models (Linderoth et al. 2006) but, to our knowledge, not yet used in ABMs in combination with a test for the number of replicates. We ran our model with 16, 32, 64, 128, 264, 256, 512, 1024, and 2048 LHS combinations and examined the results of each with Partial Rank Correlation Coefficients (PRCC) using the *pcc* function in the R package *sensitivity* (Iooss 2018). PRCCs allow the examination of parameter and response, correcting for the variation in other parameters being perturbed, making it a good tool for examining model output generated via LHS (Marino et al. 2008), and allowing us to determine how many LHS combinations were required to stabilize the results. To test if values highlighted by the PRCC analyses were actually the most influential, we calculated a z-statistic and performed pairwise comparisons following the method outlined in Marino et al. (2008).

To examine the interaction between the number of bees per 1,000 flowers and the proportion of female to male flowers, we ran an additional set of models at the full combination of bee densities (1 – 20 bees / 1,000 flowers, by 1) and proportion of female to male flowers (0.05 - 0.75, by 0.05); the remaining parameters were held at base values. Thirty-two runs were done at each parameter combination, for a total of 9,600 additional model runs.

## 3 Results

### 3.1 Sensitivity Analysis

Variation between runs decreased sharply as replicates increased (Figure 5). Based on these results, we determined that 32 replicates per parameter combination was sufficient. The parameter sensitivity analysis was then run with multiple Latin hypercube sampling (LHS) combinations.

**Figure 5:**
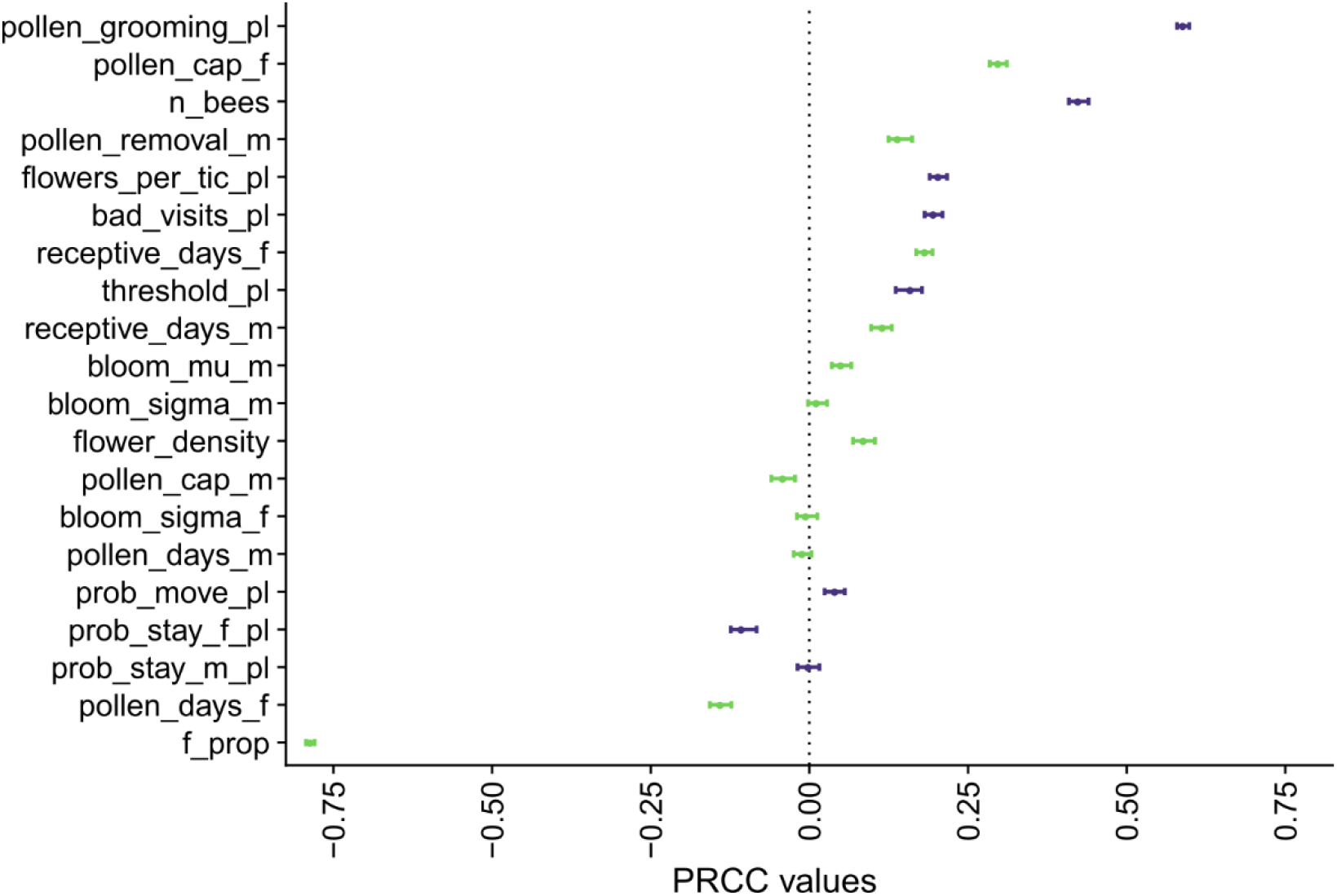
Results of PRCC-based sensitivity analysis for parameters in the agent-based model with the number of fruit produced as the response variable. Results presented are for the ‘stripmale’ orchard layout at 2048 Latin hypercube sampling combinations. Dots represent the PRCC estimate, lines the 95% CI. Green bars are plant-related parameters, purple bars are insect-related parameters.

As LHS sampling increased, the difference in PRCC values between model runs decreased—dropping from upwards of 150% down to sub-50% change between model runs with increasing samples. There was a sharp change in the output after 256 samples (Figure 6), with values above that giving more consistent results.

**Figure 6:**
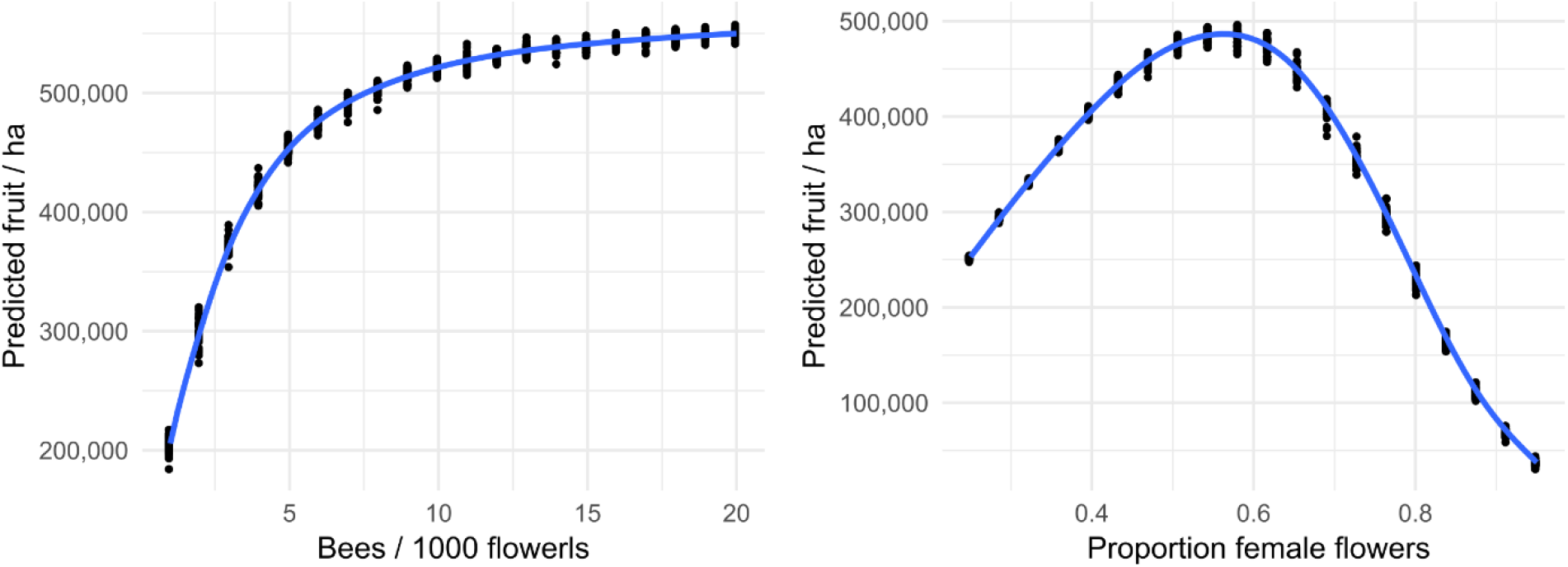
Relationship between predicted fruitset, the number of bees per 1000 flowers, and the proportion of flowers in the orchard that are female. A proportion of 0 corresponds to an all-male orchard, and a proportion of 1 corresponds to an all-female orchard. Each data point represents one of 32 simulations at the given x value, with other parameters at base values.

Interestingly, the importance of different parameters varied between Latin hypercube samples, with some parameters changing from influential to not between model runs. As there is no hard and fast rule for the number of LHS samples to sufficiently represent the parameter space and we observed potentially spurious statistical significance at higher LHS due to decreasing confidence intervals (statistical significance does not necessarily imply biological significance), we did not simply report the values for the highest LHS combination. To give a better representation of our findings, we report whether or not each predictor variable was significant in the final three Latin hypercube samples together for each landscape type (yielding a number out of 12 total model runs).

### 3.2 Plant biology

Using the base parameters, we examined the model behavior of pollen deposition over time; the trend was an increasing spread of pollen deposition as flower number increased, tapering off near the end of bloom. Our analysis identified that the density of flowers was the single most important factor for fruit production, an expected result as more flowers means more potential fruit. The proportion of female flowers (*f_prop*) was a close second, with both ∼2-3 times as influential as the next most important parameter across all four orchard types for LHS samples 512-2048 (12/12; Figure 7).

**Figure 7:**
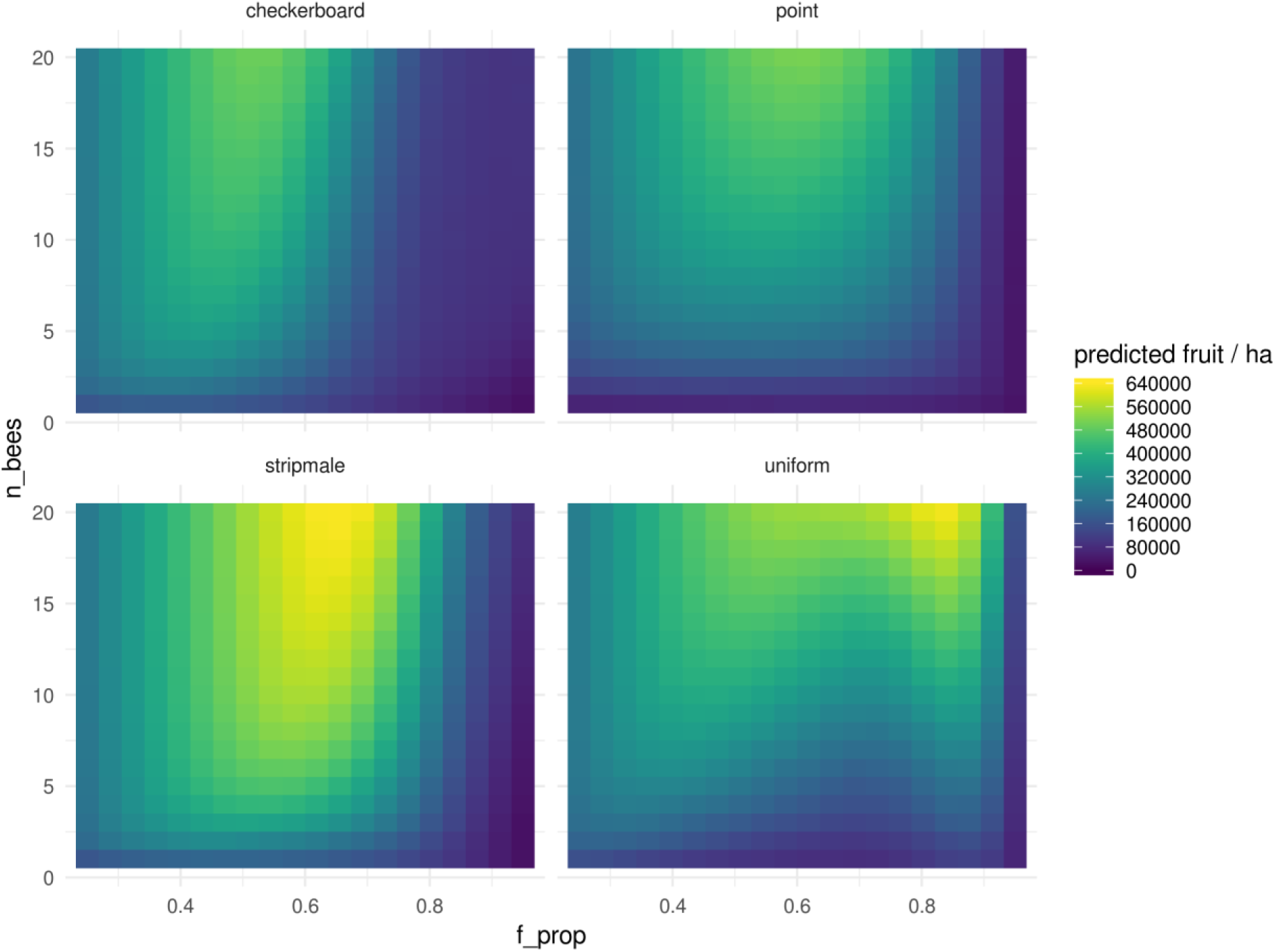
Predicted fruit set across the four orchard types and parameter values. Each pixel represents the mean fruitset for 32 model runs at that parameter combination. A f_prop of 0 corresponds to an all-male orchard, and a f_prop of 1 corresponds to an all-female orchard. n_bees represents the number of bees per 1,000 open flowers.

Much less influential, but also consistent across the four orchard types, was the amount of pollen each female and male flower contained, with more pollen on female flowers (*pollen_cap_f*) being positively correlated with fruit production, and more pollen on male flowers (*pollen_cap_m*) being negatively correlated with fruit set. Finally, with about half of the influence of pollen quantity, was the spread of flowering around the mean, with a wider distribution of female flowers (*bloom_sigma_f*) positively correlated with fruit production, and a wider distribution of male flowers (*bloom_sigma_m*) negatively correlated with fruit production in most of the orchard layouts at LHS combinations 512, 1024 and 2048. We also found that the amount of pollen produced by female flowers (*pollen_cap_f*) was somewhat influential (ranking 4^th^-10^th^ place) and positively correlated with the number of fully pollinated fruit, while the amount of pollen produced by male (ranking 4^th^-11^th^ place) was negatively correlated with the number of fully pollinated fruit.

When varying values of *f_prop* at base values for other parameters, we found that there appeared to be an optimum proportion of 50-60% female flowers in the orchard (Figure 6).

### 3.3 Pollinator biology

The most influential pollinator-related factors had 1/2 to 1/3 the influence of the strongest plant-related factors. Four factors emerged across all four orchard layouts: the number of bees per 1,000 flowers (*n_bees*), the number of flowers examined per timestep (*flowers_per_tic_pl*), the proportion of pollen removed from a flower during a flower visit (*pollen_removal_pl*), and the efficiency of grooming pollen off the body (*pollen_grooming_pl*; Figure 7). The most influential of these three was *n_bees*, which placed third in 7/12 orchard & LHS combinations and tied for third in a subsequent 5/12. The second most consistently influential pollinator parameter was *flowers_per_tic_pl*, ranking between 3^rd^ and 6^th^. The remaining two (*pollen_removal_pl* and *pollen_grooming_pl*) ranked between 3^rd^ and 8^th^. Rankings for all other variables were inconsistent. We found that increasing bee numbers at base *f_prop* values had a saturating relationship, with the most benefit gained up to 5-7 bees per 1,000 flowers (Figure 6).

### 3.4 Orchard layout

Some of the responses to changes in parameter space varied with orchard type. The ‘point’ and ‘uniform’ orchard layout benefitted nearly twice as much from an increased number of bees per 1,000 open flowers as the other two orchard types. Additionally, many parameters had a stronger correlation to fruit production in the ‘point’ orchard layout than other orchard types, including tolerance to successive low-quality flowers (*bad_visits_pl*), probability of staying on female flowers (*prob_stay_f_pl*), and the longevity of male flowers (*receptive_days_m*; Supplement).

Considering the interaction of *f_prop* and *n_bees* together across the four orchard types, we found that the ‘stripmale’ orchard outperformed all other orchard types at typical bee densities and sex ratios (Figure 9). This is surprising as the naïve assumption is that uniform orchards would produce the highest yields, but our model predicts that the ‘stripmale’ orchard layout result 40-110% higher yield, driven by bees in the flower-rich male-dominated cells becoming heavily loaded with male pollen, which they then distribute to the nearby female flowers. In contrast, bees in the ‘uniform’ orchard layout do not switch floral preference as often, as flowers of the preferred sex are available in each cell.

## 4 Discussion

### 4.1 Plant biology

Across all four orchard configurations, we found that plant-related parameters were more influential than pollinator-based parameters for both pollen deposited per flower and the number of fruit produced per hectare. For fruit production, the flower density was the most important, followed closely by the proportion of flowers that were female, with an optimal field being 50-60% female flowers. This value closely approximates values observed in modern kiwifruit orchards, where male vines produce between 3.8 – 10x more flowers than female vines (Testolin 1991; Vaissière et al. 1996) which are planted at 1/3 to 1/8 the rate of female vines (Goodwin 2012). However, the actual ratio of flowers in the field is variable (Broussard, pers. obs.). This finding, if confirmed by empirical tests, has significant implications for kiwifruit growers, by suggesting that vine planting, pruning, and flower thinning may be a better strategy to achieve sufficient pollination rather than increasing the honey bee stocking rate above the current high levels (Donovan & Read 1992), increasing male cultivar quality (Brundell 1975; González et al. 1994), or applying artificial pollination (Hopping & Hacking 1983).

Of additional significance is the fact that the amount of pollen produced by the female cultivar was positively correlated with fruit production in our sensitivity analysis. Previous studies have assumed that the quantity of pollen produced by male cultivars was the critical factor (Hopping 1984; Goodwin 1987; González et al. 1994), as bees leave the orchard in the afternoon because the majority of the pollen has been removed (Goodwin 1995). It was not until the late 1990’s that female flower pollen production was even assessed (Howpage 1999), when it was estimated to be approximately half that of common male cultivars. Since pollen is the only reward for nectarless kiwifruit flowers, this finding is intuitive. In a scenario where 50% of the flowers are female, but they each have less pollen than male flowers, they may run out of pollen before male flowers do, limiting the pollination window. This may also explain the reason that the quantity of male pollen per flower was negatively correlated with fruit set.

### 4.2 Pollinator biology

Three insect-related parameters surfaced as influential factors in our model. The first, the number of bees per 1,000 flowers, is well-attested in the kiwifruit literature (Palmer-Jones et al. 1976; Clinch 1984; Donovan & Read 1992; Pomeroy & Fisher 2002; Goodwin et al. 2017). Being a measure of insect visitation, which has been identified as a key component of pollination in cropping systems as well as natural areas, this is not surprising. However, visitation on its own has been shown to be a poor predictor of pollination due to the variable nature of pollinators (King et al. 2013). While we do not have single-visit deposition as a parameter in our model—it arises organically based on the pollen available on the insect body—two factors related to this value were significant across all orchard types: the proportion of pollen removed from a flower and the proportion of pollen on the body transferred to flowers. We chose a wide range for these parameters, as there is little baseline information; further study of these factors would be useful to inform these parameter ranges.

An orchardist may be able to increase honey bee density through higher stocking rates (though see (Clinch 1984; Rollin & Garibaldi 2019), but the other factors cannot be directly controlled, though they do vary between insect species. This result could be tested empirically by bringing in other managed pollinators or increasing numbers of local pollinators, though there is suggestive evidence that multiple pollinator species do have a positive effect through complementarity (Garibaldi et al. 2013; Broussard et al. 2022). Given that pollinator-related factors are more difficult for growers to manage, it is serendipitous that they appear to have significantly less influence than plant-based metrics, which can be much more precisely controlled through planting and pruning.

### 4.3 Orchard layout

Orchard layout has previously been shown to affect bee behavior in kiwifruit orchards, with Jay & Jay (1984) reporting early on that having the canopy overhead on a pergola system caused more bees to cross between rows than a row-based trellis system. We found that changes in orchard layout resulted in up to a factor of two difference in fruit production at typical bee densities and flower sex ratios.

We also found that the ‘point’ orchard layout, with the longest distance to travel between male and female flowers, showed the worst performance in total fruit production. This is corroborated in the literature, where increasing distance from male vines causes declines in fruit set and seed number (Testolin 1991; Goodwin et al. 2013). Interestingly, the ‘point’ orchard outperforms the ‘uniform’ orchard at low bee densities when female flowers comprise 60-80% of all flowers, driven by bees loaded with pollen exiting high-flower density male area and depositing pollen on nearby flowers.

Our simulations indicate that the ‘stripmale’ layout, which has been increasing in popularity in New Zealand for some decades (Goodwin, pers. comm.) may outperform the hypothetically optimal ‘uniform’ configuration at all but extremely high bee densities and low male flower numbers. The good performance of this configuration could explain its popularity, and is a promising result for trialing novel orchard layouts.

### 4.4 Conclusion

Pollination involves complex interactions between plants and pollinators, and variation in plant or pollinator biology can lead to variability in pollination services that are difficult to predict. Cross-pollination in a functionally dioecious system is critical for fruit and seed set, yet has not been adequately modelled to date. Our model of kiwifruit can serve as a test case for modelling this challenging pollination system. Several important trends emerged from our model, including the prediction that plant traits may be more influential than pollinator traits for predicting pollen deposition and fruit yield. This suggests that simple and inexpensive orchard management practices may be more effective than costly hive overstocking or artificial pollination. Although our model agrees with findings from the literature where they exist, it is not fully validated through empirical study. Collecting data on bee foraging rates across numerous orchard types would be prohibitively expensive, however validation could be achieved with cameras and automatic bee monitoring combined by full-orchard yield data. Should the model continue to align with field observations, there is strong potential to use it to test a multitude of novel planting regimes, too costly to conduct in field conditions, and identify those suitable for trial on-orchard. Additionally, there is significant scope to expand this model to examine synergistic effects between honey bees and other pollinators, or a combination of pollination strategies, and to extend it to other cropping and natural ecosystems.

## 4.5 Acknowledgements

We would like to thank R. Mark Goodwin for his wealth of knowledge of kiwifruit pollination and direction toward datasets which could be used to parameterize the model. We would also like to thank Paul Martinsen, Warrick Nelson and Flore Mas for reviewing an early draft of this manuscript and two anonymous reviewers for their helpful feedback.

